# Elevated phosphorylated α-synuclein and phosphorylated tau in Lrrk2 p.G2019S-mutant mice following influenza A pneumonitis

**DOI:** 10.1101/2025.07.31.667659

**Authors:** Michaela O. Lunn, Karan Thakur, Nathalie A. Lengacher, Quinton Hake-Volling, aSCENT-PD Investigators, Julianna J. Tomlinson, Subash Sad, Earl G. Brown, Michael G. Schlossmacher

## Abstract

**Background:** Influenza virus exposure elevates the incidence of parkinsonism. We and others previously discovered that allelic variants at the Parkinson’s-linked *LRRK2* locus modulate host responses to virulent microbes.

**Objective:** We asked whether *Lrrk2* mutations modify disease outcomes in adult animals following a nasally acquired lung infection.

**Methods:** We inoculated C57BL/6J mice of mutant knock-in *Lrrk2* genotypes with influenza A virus, H1N1-serotype (1 x LD_50_ = 2,000 plaque-forming units).

**Results:** During H1N1-induced pneumonitis neither homozygous nor heterozygous mutations of kinase activity-increasing *Lrrk2^G2019S^* or kinase-dead *Lrrk2^D1994S^* altered the course of sickness in mice when compared to wild-type littermates, as determined by survival rates, viral titres and weight changes in both sexes. However, six weeks after inoculation brains of H1N1-exposed, homozygous Lrrk2 p.G2019S animals showed higher pSer_129_ α-synuclein and pSer_199_ tau levels than mock-treated, mutant mice (*P<0.01*). The ratios of phosphorylated tau-to-total tau and phosphorylated α-synuclein-to-total α-synuclein also rose in H1N1-exposed Lrrk2 p.G2019S mice (*P<0.05*). Further, we found that nitrotyrosination of the brain proteome was significantly increased in *Lrrk2^G2019S^*survivors vs. mock-exposed littermates (*P<0.05*). Brain H_2_O_2_ concentrations were elevated in male, H1N1-inoculated wild-type animals (with a trend seen in females and *Lrrk2^G2019S^* mice), an effect that was abrogated in kinase-dead *Lrrk2^D1994S^* mice.

**Conclusion:** Homozygous *Lrrk2^G2019S^*-mutant mice that survived infection by a life-threatening, pneumotropic RNA virus show changes in brain levels of oxidative stress, pSer_199_ tau and pSer_129_ α-synuclein. These results could be of relevance to the initiation of neurodegeneration-linked changes in humans and may help explain differences in the penetrance rate and expressivity of *LRRK2* mutants.

## INTRODUCTION

Parkinson’s disease (PD), a complex neurodegenerative disorder, is increasing in prevalence, yet the etiology of the disorder remains elusive. It is thought that PD is caused by a complex interplay between genetic risk, environmental triggers, and ageing (1). Several environmental risk factors have been identified, including non-neurotropic viral infections (2,3). For example, the 1918 influenza pandemic, caused by an influenza A serotype H1N1 variant, was associated with the subsequent emergence of von Economo’s encephalitis, including a form linked to post-infectious parkinsonism (4,5). More recently, a positive influenza exposure history (up to fifteen years prior) has been linked to a rise in PD incidence (6). In mice, infection following nasal inoculation with the highly pathogenic avian influenza serotype H5N1 (a neurotropic virus) led to α-synuclein phosphorylation and aggregation as well as dopamine cell loss in the brain of wild-type mice (7,8). Furthermore, H1N1 has been shown to increase α-synuclein levels *ex vivo*, which was ameliorated by pre-treatment with an anti-influenza drug (9).

Beyond environmental risk factors, variants at >90 loci have been identified and attributed to altered PD risk, including mutations in the *LRRK2* gene, which account for ∼2% of cases of parkinsonism seen in specialty clinics (10–12). Mutations in this gene confer risk for both familial and sporadic forms of PD, although the mechanisms remain elusive (13,14). Notably, there is incomplete penetrance associated with *LRRK2* mutations, including for the commonest variant, p.G2019S (15). This suggested to us that disease manifestation in *LRRK2* mutation carriers may require a second trigger, either genetic or environmental; we and others have postulated that the latter factor could be microbial in nature (16–21).

Importantly, mutations at the *LRRK2* locus have also been linked to Crohn’s disease, leprosy, and systemic lupus erythematosus (22–24). Each of these diseases is characterized by chronically dysregulated inflammation. The shared risk locus among four distinct diseases raises the issue of whether an evolutionarily preserved function of LRRK2 involves regulation of immune responses. LRRK2 is expressed at higher levels in the periphery (*e.g.,* bone marrow, lung) than the brain, and is more abundant in cells of the immune system (neutrophils, monocytes, and macrophages) than neurons (25,26). Further, the protein can be upregulated by infectious pathogens and inflammatory mediators (*e.g.,* IFN-γ and NF-κB), suggesting that LRRK2 is involved in pathogen defence and immune system responses (26–29). Several studies have demonstrated effects by Lrrk2 on bacterial pathogenicity in a microbe-dependent manner (18,23,30,31). We previously also investigated the role of Lrrk2 in response to viral infections; there, we demonstrated that disease outcomes following nasal inoculation with reovirus [isotype type-3 Dearing (T3D)] in mouse pups were affected by allelic variants of *Lrrk2* in a sex-dependent manner (18).

Here, we tested whether allelic variants of *Lrrk2* could alter the course of a potentially lethal lung infection in adult C57BL/6J mice and reveal any brain pathology following exposure to a pneumotropic RNA virus. We hypothesized that a kinase-hyperactive variant of *Lrrk2* may lower viral replication in the lung and alter survival rates in adult animals. Further, we postulated that this infection may lead to changes in neurochemical markers in surviving animals. To test this, we employed a mouse-adapted influenza A virus, serotype H1N1 (A/Fort Monmouth/1/1947), to induce pneumonitis in wild-type and *Lrrk2* knock-in mutant mice. We found that neither the *Lrrk2* p.G2019S nor p.D1994S mutant altered acute disease outcomes. However, having survived influenza H1N1 infection led to increased levels of oxidative stress and changes in the metabolism of α-synuclein and tau in the brains of homozygous *Lrrk2* p.G2019S animals when compared to their littermates.

## METHODS

### Mouse Colonies

All mouse colonies were maintained on a C57BL/6J background using heterozygous breeding pairs. Animal experiments were performed under Canadian Councils on Animal Care guidelines and approved by the Ethics Board of the Animal Care Committee at the University of Ottawa. Experiments involving viral infections were done in a containment level 2 biohazard animal facility. *Lrrk2* p.G2019S (MGI:5295368) and *Lrrk2* p.D1994S (MGI:5295371) knock-in mice were kindly provided by Dr. Derya Shimshek, as described (32). Both female and male mice were used in this study, as indicated.

### Nasal Infections with Mouse-Adapted Influenza A H1N1 Virus

Generation of the mouse-adapted variant of influenza A H1N1 [A/Fort Monmouth/1/1947 (‘FM-MA’)] was previously described (33). Adult mice aged 5-7 weeks were anesthetized with 3% isoflurane in O₂ at 1 L/min for 1 minute. A dose of 2×10^3^ plaque-forming units (PFU) of FM-MA H1N1 (prepared in 50µL of PBS) was administered to the nostrils, inducing severe pneumonitis at ∼50% lethality (1 LD_50_). Mice were observed for 14 days post-inoculation (dpi) and weighed twice daily from 4-14 dpi. Endpoint criteria to prompt euthanasia included a moribund state, labored breathing, or ≥30% body weight loss. Mice that survived the acute viral infection (evident by survival at 14 dpi) were maintained for 4 additional weeks to a total of 6 weeks post-inoculation (wpi) (**Fig. 1a**). In parallel to H1N1-exposed mice, mock-inoculated animals received 50µL of PBS without virus.

**Figure 1.**
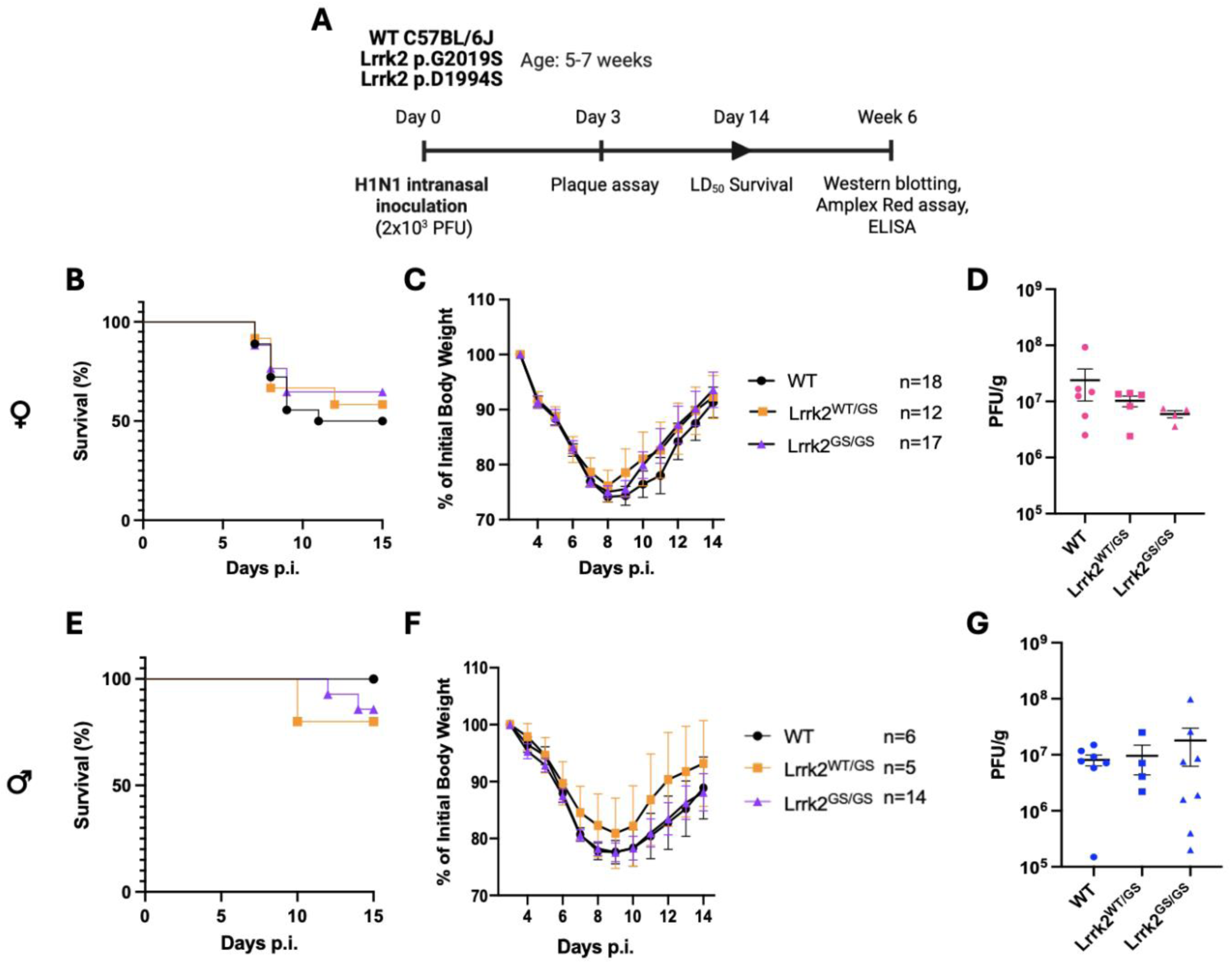
*Lrrk2^G2019S^*alleles do not alter survival, weight change, or viral titres in the lungs of mice nasally inoculated with influenza A virus, H1N1. Experimental timeline employed in this study (**A**). Adult mice (males and females; aged 5-7 weeks) were nasally inoculated with 2 x 10^3^ plaque-forming units (PFU) of influenza A virus (serotype H1N1). Survival is graphically displayed as percent survival against days post inoculation (p.i.) (**B, E**), and weights are shown as percent of initial body weight against days p.i. (**C, F**). Data are shown separately for female (**B-D**) and male mice (**E-G**). Viral titres were measured in the lungs at 3 days p.i (**D, G**). Each symbol represents one animal. Error bars represent mean ± SEM. WT = wild-type; GS = *Lrrk2^G2019S^* mutant allele. Statistical tests used: log rank (Mantel-Cox) test for survival curves; one-way ANOVA for viral titre analyses.

### Influenza A Virus Serotype H1N1 Plaque Assay

Mice that had been nasally infected with 2×10^3^ PFU of H1N1 were sacrificed at 3dpi. Lungs were snap frozen, then lysed in PBS (3:1, v:w) using metal beads in a Magna Lyser (Roche) at maximum speed for 10 seconds, then freeze-thawed once. MDCK cells were grown in 6-well plates to 90% confluency. Lung homogenates were centrifuged (5,000 RPM) and supernatants were serially diluted (10-fold) in serum-free DMEM. MDCK monolayers were washed twice with PBS, overlaid with 100µL of each dilution, and incubated at 37 °C, 5% CO₂ for 30 minutes, with rocking at 15 minutes. After one hour, cells were overlaid with 3 mL of 1:1 1.2% agarose:2xDMEM containing 0.1 µg/mL TPCK-trypsin and incubated for 3 days. Cells were fixed in methanol:acetic acid (3:1) for one hour. Overlays were removed under tap water and stained with Coomassie blue for 30 minutes. Plaques were counted visually. Viral titres are reported as PFU per gram of tissue.

### Protein Extraction

Protein was extracted by tissue homogenization, as described above for viral titre analyses. Aliquots were prepared in 1% Triton-X 100 and 2x protease with phosphatase inhibitors (Thermo Scientific) and centrifuged at 14,800 RPM for 30 minutes at 4°C. Protein concentrations were determined using a bicinchoninic acid (BCA) assay kit (Thermo Fisher Scientific, 23227), according to the microplate procedure.

### ELISA Quantification of α-Synuclein and Tau

Lysates were measured for total α-synuclein using a sandwich ELISA, as described previously (34,35). α-Synuclein protein concentrations were normalized to the total protein concentration in each sample. Total murine tau protein was measured using the Tau (Total) Mouse ELISA Kit (KMB7011, Invitrogen). Mouse phospho-tau (phospho-serine 199) abundance was measured using the Mouse Tau (Phospho) [Ser199] ELISA Kit (KMB7041, Invitrogen).

### Western Blotting

Protein samples (40µg per lane; 75µg for anti-nitrotyrosine) were mixed with 1xNuPAGE™ LDS buffer, boiled for 5 mins at 95 °C, and run on 4-12% gradient SDS-MES gels under reducing (non-reducing for anti-nitrotyrosine) conditions. Proteins were transferred to PVDF membranes by semi-dry transfer in TRIS-glycine-methanol buffer. Anti-nitrotyrosine antibody (Cell Signalling, 9691S) blots were imaged on film, quantified via ImageJ, and normalized to Ponceau S staining. Phospho-α-synuclein (Ser129) (Abcam, ab51253), hSA4 (a non-commercial, polyclonal Ab, that was raised and affinity-purified against recombinant, full length, human α-synuclein as described (Mollenhauer et al. <u>2008</u>)) and Plk2 (Novus Biologicals, NBP3-04961) antibody blots were imaged using a BioRad ChemiDoc and quantified using ImageLab. Signals for phosphorylated α-synuclein and total α-synuclein were normalized to actin (Santa Cruz, sc-47778) as a loading control (Plk2 was normalized to Ponceau S staining) and further normalized to internal blot controls for inter-blot quantification.

### Quantification of Reactive Oxygen Species

For H_2_O_2_ quantification in tissues, the Amplex® Red hydrogen peroxide/peroxidase assay kit (Invitrogen, A22188) was used to monitor endogenous levels of H_2_O_2_. Sagittally cut hemibrains were lysed in PBS, as described above. Levels of H_2_O_2_ in homogenates were measured per manufacturer’s instructions. Obtained H_2_O_2_ values (μM) were normalized to tissue weight (g).

### Statistical Analyses

All statistical analyses were performed using GraphPad Prism version 11 (GraphPad Software, San Diego, CA, USA). Differences between groups in survival assays were assessed using the log-rank (Mantel-Cox) test. Differences between groups were assessed using a two-way ANOVA with an uncorrected Fisher’s LSD post-hoc test. Data are shown as mean ± SEM where applicable and as described in the figure legends. Data are displayed with *P* values represented as \**P*<0.05, *\*\*P<*0.01, \*\*\**P*<0.001, \*\*\*\**P*<0.0001.

### Data, Protocols, and Materials Availability

The datasets and images used and/or analyzed in this study, along with a Key Resources Table (KRT) containing detailed protocols and key lab materials along with their persistent identifiers, are available on Zenodo: https://doi.org/10.5281/zenodo.16576378.

## RESULTS and DISCUSSION

### Lrrk2 p.G2019S mice show the same disease course as wild-type mice during H1N1 pneumonitis

We first investigated the role of enhanced Lrrk2 kinase activity in host responses using a non-neurotropic virus in *Lrrk2^G2019S^* and wild-type littermates (**Fig. 1**). Influenza A virus, serotype H1N1 (FM-MA strain), has lung tropism that can cause fatal pneumonitis in mice at specific doses. Notably, Lrrk2 is highly expressed in the lungs (18,32). Adult mice aged 5-7 weeks were nasally inoculated with a previously established LD_50_ dose (2×10^3^ PFU) (33); animals were either sacrificed at 3dpi for viral titre analysis in their lungs, or assessed for survival with survivors then being aged (**Fig. 1a**). Survival rates and weights were measured up to 14dpi in female (**Fig. 1b, c**) and male (**Fig. 1e, f**) animals.

Female mice in general showed higher mortality rates than males, which is consistent with previous reports using various strains of H1N1 in C57BL/6J mice (36,37). However, we did not identify any differences in the survival of Lrrk2 p.G2019S knock-in mice compared to wild-type littermates. Similarly, there were no significant differences between genotypes (and mutant allele numbers) in initial weight loss or subsequent weight gain, which served as surrogates for sickness and recovery, respectively. As a third outcome of H1N1 infection, we measured viral titres in the lungs at the peak of infection, namely at 3dpi, in a separate cohort. Although we observed a trend for lower titres in female *Lrrk2^G2019S^* homozygous mice (which we had expected from our reovirus T3D results in pups) (18), greater variability in titres was seen in the corresponding males. Overall, no significant differences were calculated for viral titres at 3dpi between the genotypes (**Fig. 1d, g**).

### Lrrk2 kinase activity is not required for acute host defence against H1N1 infection

Because we did not identify any modulation of survival conferred by the hyperactive kinase p.G2019S mutant, we next investigated whether the loss of Lrrk2 activity would affect disease outcomes of H1N1 infection, as we had seen in response to reovirus-T3D in newborns (38). Using the same approach as above, we inoculated mice carrying the Lrrk2 kinase-dead p.D1994S knock-in mutation. We found that both male and female *Lrrk2^D1994S^* homozygous and heterozygous mice showed the same survival rate and weight changes as wild-type mice following H1N1 infection (**Suppl. Fig. 1a, b, d, e**). To probe further, we measured H1N1 viral titres in the lungs of these mice at 3dpi and again did not identify any group differences between wild-type, heterozygous, and homozygous Lrrk2 p.D1994S animals (**Suppl. Figs. 1c, f**).

These collective results suggested to us that the Lrrk2 kinase activity itself did not show a detectable effect in the response of adult mice during the acute phase of H1N1 pneumonitis.

### H1N1 survivors do not show evidence of viral protein expression in the brain

We next probed for post-infection impacts of H1N1 virus exposure on the brain in surviving mice and any effect potentially attributable to the *Lrrk2^G2019S^* knock-in mutation. Homozygous mice that survived the H1N1 infection were first aged for four additional weeks to a total of 6wpi (**Fig. 1a**). We then examined homogenates from H1N1- and mock-inoculated animals with an anti-H1N1 immune serum from rabbits by Western blotting. Using an infected lung at 3dpi as a positive control, we did not detect any virally encoded H1N1 proteins, *i.e.,* nucleoprotein NP and matrix protein M1, in the brains of our cohort of surviving mice (data not shown), suggesting that H1N1 was not detectable in the brain under these conditions, as expected.

### Phosphorylated tau is increased in Lrrk2 p.G2019S mutant brain following H1N1 infection

Brain autopsies previously performed on post-encephalitic cases of parkinsonism in victims of the influenza A (H1N1) virus pandemic revealed evidence of a tauopathy (39). Of note, the observed brain pathology in *LRRK2-*linked cases of PD is variable and includes hallmark α-synuclein aggregates, phosphorylated tau-linked pathology, neuronal loss without inclusion formation, or mixed pathologies (10,40,41). Therefore, we first measured the abundance of total tau followed by the level of phosphorylated tau (pSer_199_ tau) in the soluble fraction of total brain homogenates by ELISA from H1N1-infected survivors vs. mock-treated animals at 6wpi (**Fig. 2a-c**). When measuring total tau in these lysates, we observed no difference in an infection- or mutation- (**Fig. 2a**) or sex-dependent manner (**Suppl. Figs. 2a, 3a**). When quantifying pSer_199_ tau, two-way ANOVA revealed a significant infection x genotype interaction (*F*(1, 23)=5.418, *P=0.0291*; **Fig. 2b**). There, pSer_199_ tau was significantly increased in infected Lrrk2 p.G2019S mice compared to mock-treated, mutant mice (*P=*0.0099), an effect that was absent in wild-type mice. We next calculated the ratio of phosphorylated-to-total-tau protein in these samples (**Fig. 2c**), where we identified a significant rise in infected, *Lrrk2* p.G2019S-mutant animals compared to mock-treated mutant mice. H1N1-infected, mutant mice also showed a significantly higher ratio vs. infected wild-type mice (*P<0.05*; **Fig. 2c**). Two-way ANOVA revealed a trend towards significance for the latter ratio (*P=0.0521*).

**Figure 2.**
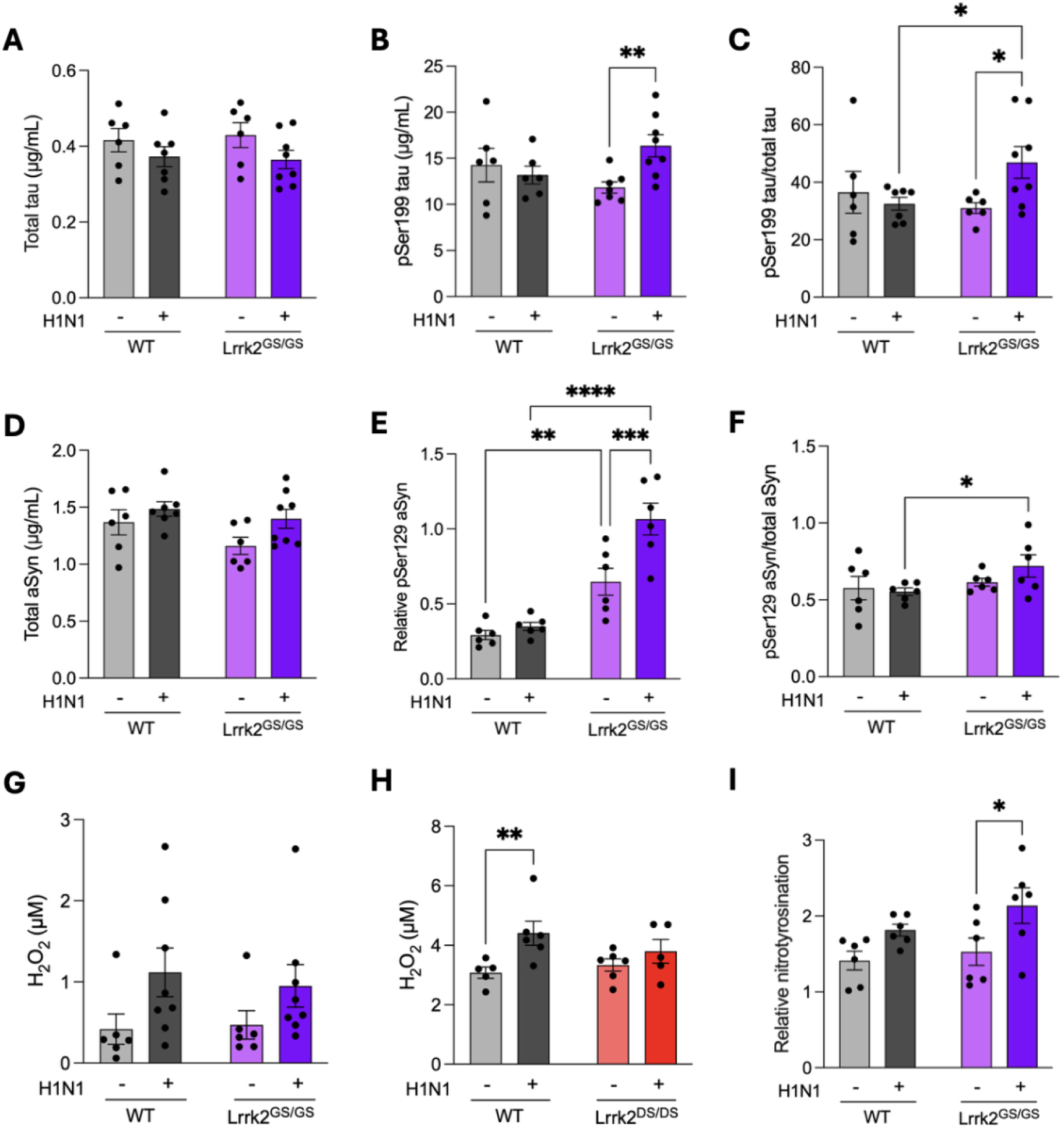
Lrrk2 p.G2019S expression alters tau and α-synuclein metabolism as well as oxidative stress in the brains of mice following H1N1 influenza pneumonitis. Adult mock-(H1N1 -) and H1N1-infected (H1N1 +) mice (males and females are shown; ages, 5-7 weeks) that survived past 14 days post-inoculation were aged for an additional 4 weeks and their brains were collected. Bar graphs indicate levels of total tau protein (**A**), phosphorylated tau (pSer_199_) (**B**), the ratio of phosphorylated tau-to total-tau levels (**C**); total α-synuclein (performed by ELISA; **D**), phosphorylated α-synuclein (pSer_129_) (**E**), and the ratio of phosphorylated α-synuclein-to-total α-synuclein (both performed by Western blotting; **F**). Levels of H_2_O_2_ in two separate cohorts of mice (**G, H**). Protein nitrotyrosination content of the brain (**I**) (H_2_O_2_) and nitrotyrosination measurements shown are relative to brain weight). Each symbol represents one animal. Error bars represent mean ± SEM. WT = wild-type; GS = Lrrk2 p.G2019S mutant mice; DS = Lrrk2 p.D1994S mutant mice. Statistical analysis: two-way ANOVA with uncorrected Fisher’s LSD post-hoc test. *P*-value <0.05 (*), *P*-value<0.01 (**), *P*-value <0.001 (***), *P*-value<0.0001 (****).

These findings indicated that soluble pSer_199_ tau concentrations were elevated in the brain following H1N1 infection, which was linked to the expression of a homozygous *Lrrk2^G2019S^* knock-in mutation. When probing the same homogenates for GSK3β, a cognate kinase for pSer_199_ tau (both active and inactive forms were examined by Western blotting), results did not reveal any infection-, sex- or mutation-dependent effect (data not shown). To date, we have not yet examined other tau kinases.

### Phosphorylated α-synuclein is increased in Lrrk2 p.G2019S mutant brain following H1N1 infection

We next examined the metabolism of α-synuclein in the same animals (**Fig. 2d-f**). We had previously seen a *Lrrk2^G2019S^*-dependent rise in total α-synuclein in the brains of reovirus (T3D)-infected newborn mice (18). We therefore performed Western blotting and ELISA of total brain homogenates from surviving females and males, probing for levels of total synuclein and pSer_129_ α-synuclein. Two-way ANOVA revealed an effect of the preceding H1N1 infection on total α-synuclein in the brains of mutant females (*F*(1, 8)=7.32, *P=*0.0268 **Suppl. Fig. 3d**), but not in males (**Suppl. Fig. 2d**); nonetheless, when sexes were pooled the infection effect remained significant (*P=0.0474*) (**Fig. 2d**). Further, we detected a rise in pSer_129_ α-synuclein levels in surviving animals (**Fig. 2e**). Two-way ANOVA revealed a significant genotype effect (*F*(1, 20)=54.73, *P<0.0001*), a significant infection effect (*F*(1, 20)=10.76, *P=0.0037*) and a significant genotype x infection interaction (*F*(1, 20)=6.186, *P=0.0218*), which was largely driven by female animals (**Suppl. Fig. 3e**). Intriguingly, female Lrrk2 p.G2019S mice showed elevated pSer_129_ α-synuclein in the brain at baseline (**Suppl. Fig. 3e**). When we calculated the ratio of phosphorylated-to-total α-synuclein levels, we identified a higher ratio in infected mutant mice compared to infected wild-type animals by pairwise comparisons (*P<0.05*) (**Fig. 2f**), which was seen foremost in males (**Suppl. Fig. 2f**); two-way ANOVA revealed significance for this interaction (*P=0.0381*).

These results suggested to us that H1N1 pneumonitis can lead to dysregulation of soluble, phosphorylated α-synuclein in the brain of surviving homozygous Lrrk2 p.G2019S animals. Given that polo-like kinase-2 (Plk2) has been identified by several teams, including ours, as a cognate enzyme for phosphorylation of α-synuclein at Ser_129_ (42–44), we also probed for it in brain lysates by Western blotting. We observed a significant rise in Plk2 protein in the brain of mutant females at baseline (*P<0.05*) (**Suppl. Fig. 3g**), which was worsened by a preceding H1N1 infection in wild-type animals; this effect was not seen when sexes were pooled (data not shown). Future work will determine whether Plk2 enzyme activity itself (or dephosphorylation of pSer_129_ α-synuclein) was altered in the brains of animals that survived H1N1 pneumonitis.

### H1N1 pneumonitis leads to elevated oxidative stress in the brains of Lrrk2 p.G2019S mice

We next measured the levels of oxidative stress in the brains of surviving animals of three distinct homozygous genotypes (**Fig. 2g, h**) in two cohort comparisons. The concentration of H_2_O_2_ in homogenates from H1N1-infected surviving mice was quantified using the Amplex® Red assay. There, we identified an effect of pneumonitis on H_2_O_2_ levels in a cohort of wild-type vs. *Lrrk2* p.G2019S animals using two-way ANOVA, namely for infected male wild-type vs. mock-treated mice (*F*(1, 12)=10.20, *P=0.0077*) (**Suppl. Fig. 2h**); however, no significant difference was found for other comparisons in this cohort (**Fig. 2g**). When measuring H_2_O_2_ concentrations in a separate cohort, where we compared wild-type to *Lrrk2* p.D1994S kinase-dead mutant mice, we again observed a significant difference in male wild-type mice (*F*(1, 18)=7.869, *P=0.0117*) (**Fig. 2h; Suppl. Fig. 1g,h**). In post-hoc analysis, preceding H1N1-infection led to significantly higher H_2_O_2_ levels in wild-type animals compared to mock-treated mice (*P=0.0087*); a trend for a rise in brain H_2_O_2_ was seen in both female and male Lrrk2 p.G2019S-mutant mice in an infection-dependent manner (**Fig. 2g; Suppl. Figs. 2h, 3h**). Intriguingly, no elevation in brain H_2_O_2_ concentrations was detected in H1N1-exposed, kinase-dead Lrrk2 p.D1994S animals (**Fig. 2h; Suppl. Fig. 1g, h**).

As a secondary measurement of oxidative stress, we investigated irreversible protein nitrotyrosination in protein extracts from the brains of the H1N1-infected mice at 6wpi (**Fig. 2i**). Using Western blotting and densitometry, we identified a significant effect of H1N1-exposure on protein nitrotyrosination in all females (*F*(1, 8)=13.34, *P*=0.0065) (**Suppl. Fig. 3i**). When analyzing corresponding levels in males, two-way ANOVA revealed significant effects for infection (*F*(1, 8)=8.484, *P*=0.0195) and genotype (*F*(1, 8)=13.62, *P*=0.0061) (**Suppl. Fig. 2i**). Post-hoc analyses confirmed that male H1N1-exposed Lrrk2 p.G2019S mice contained higher nitrotyrosination levels than mock-treated, mutant animals and inoculated, wild-type mice.

Recently, two teams—including ours—identified a role for Lrrk2 p.G2019S in upregulating NADPH oxidase-2 (NOX2) activity, which leads to a burst in ROS production by select immune cells (38,39); therefore, we also probed for the NOX2 subunit p47^phox^ within its active complex, which we previously found hyper-phosphorylated by Lrrk2 p.G2019S in neutrophils (45). To date, analysis of the same brain lysates from H1N1-infected survivors vs. mock-inoculated animals, as examined above, using anti-p47^phox^ by Western blotting, remained inconclusive (data not shown).

In sum, we have shown that adult C57BL/6J mice that survived a life-threatening pneumonitis due to a viral infection of their lower respiratory tract featured neurochemical changes in the brain. The pneumotropic H1N1 infection led to a significant increase in oxidative stress and elevation of both phosphorylated α-synuclein and phosphorylated tau in adult *Lrrk2* p.G2019S-mutant mice at 6 weeks after inoculation, *i.e.*, at a time when survivors had recovered from their acute illness. Our results demonstrate that neither the *Lrrk2* p.G2019S nor the *Lrrk2* p.D1994S mutant conferred a benefit—or elevated risk—during the course of an *acute* viral H1N1 lung infection, as evident by similar survival rates, viral titres, and weight changes (**Fig. 1**). In our previous studies, *Lrrk2* p.G2019S had conferred protection in a bacterial *Salmonella typhimurium* sepsis model (18,47) and *Listeria monocytogenes* infection model (45), but had worsened survival during reovirus (T3D)-induced encephalitis with a female sex effect (18). These results had generated a hypothesis by which the *Lrrk2* p.G2019S mutant may increase host defences against select microbial infections outside the brain but confers greater risk of neuronal loss when an infection reaches the nervous system (as compared to wild-type animals and *Lrrk2* p.D1994S mice). Informed by our previous studies, contributing factors to worse brain health included an excess in ROS production (such as via upregulated NOX2 activity), augmentation of microglial reactivity, and enhanced chemotaxis of myeloid cells into the brain (18). Because H1N1 does not have direct tropism for the brain, we postulate that pro-inflammatory responses initiated in the periphery reached the brain and affected its neuronal and/or glial metabolism in our pneumonitis model described here, a concept that matches the conclusion reached by Cadar et al. for post-encephalitic parkinsonism following the H1N1 pandemic a century ago (39).

Upregulation of α-synuclein has been reported *in vivo* in response to various inflammatory triggers (48–50). Here, we detected a *Lrrk2* p.G2019S-based effect on the pSer_129_ α-synuclein steady-state in female and male mice after H1N1 infection. Hence, our findings could be of relevance to the risk association of an influenza A virus infection with the subsequent incidence of late-onset PD (6), especially in carriers of a mutant *LRRK2* genotype that develop a synucleinopathy (41,51).

*LRRK2*-linked PD exhibits pleiotropic brain pathology (40,41,52,53), and mutations in it have also been linked to the tauopathy of progressive supranuclear palsy (PSP) (52–55). In this context, we assessed the effects of a preceding H1N1 infection on tau metabolism. We identified a significant infection and genotype interaction regarding the steady-state of pSer_199_ tau; there, levels were significantly increased in *Lrrk2* p.G2019S-mutant (but not wild-type) female mice following H1N1 infection. In humans, a tauopathy featuring insoluble neurofibrillary tangles has not only been described in cases of post-infectious parkinsonism (56), but also in the subacute sclerosing panencephalitis caused by measles virus (57–60). In future work, peripheral immune mediators, such as circulating cytokines, chemokines and infiltrating immune cells, will need to be assessed for their mechanistic contributions to neurochemical changes observed in our pneumonitis model.

Our study has limitations. One, analyses were restricted to 6 weeks post inoculation, which may not capture the full trajectory of disease resolution. Two, we did not evaluate age-dependent susceptibility, as all mice were relatively young (5-7 weeks old) and immune responses as well as the integrity of the blood-brain-barrier are known to vary across the lifespan. Three, several studies have demonstrated that tau phosphorylation is highly sensitive to ambient factors and stress conditions, such as hypothermia, fever, anesthesia, and the sleep-wake cycle (61–65). In our studies, we were attentive to these potential modifiers, but others (not yet identified) could have played a modulatory role. Four, we have not yet uncovered the mechanisms behind the hyperphosphorylation of α-synuclein and tau in *Lrrk2* p.G2019S mice that survived H1N1 pneumonitis (of note, Lrrk2 itself is not considered a neuronal kinase for tau or α-synuclein). Five, hyperphosphorylation of two proteins in soluble brain lysates does not equate the generation of insoluble, amyloidogenic aggregates associated with tangle formation and Lewy-type inclusions within neurons later in life; however, it could serve as a precursor event. Lastly, more animals added to each experimental arm could potentially have shed even more light on sex effects in our neurochemical readouts.

Nonetheless, our results suggest that infection by a pneumotropic RNA virus can augment oxidative stress in the brains of C57BL/6J wild-type mice and elevate phosphorylated α-synuclein and phosphorylated tau in *Lrrk2* p.G2019S knock-in animals six weeks after nasal inoculation despite a lack of direct neurotropism by H1N1 (and no detectable viral proteins in the brain at that time). Our findings in adult mice, housed under controlled conditions, add to the concept that interactions between select virulent microbes, including those targeting peripheral organs, and the host’s genome could serve as initiators of neurodegeneration-linked changes in adult human brain.

## AUTHOR CONTRIBUTIONS

*Study design:* M.O.L., J.J.T., E.G.B., M.G.S.; *Writing and Figure preparation:* M.O.L., K.T., J.J.T., E.G.B., M.G.S. All authors reviewed and / or edited the manuscript and approved of the submitted version. *Experiments:* M.O.L., K.T., N.L., Q.H-V., performed experiments and / or data analysis. *Analysis:* M.O.L., K.T., N.L., Q.H-V., J.J.T., S.S., E.G.B., M.G.S. performed data interpretation. *Study supervision*: J.J.T., E.G.B., M.G.S. *Overall responsibility:* M.G.S. aSCENT-PD Investigators include: Benjamin Arenkiel, Gianfilippo Coppola, Zhandong Liu, Brit Mollenhauer, Josef Penninger, Maxime Rousseaux, Natalina Salmaso, Christine Stadelmann, Michael G. Schlossmacher, Julianna J. Tomlinson, John M. Woulfe.

## ACKNOWLEDGEMENTS

We are grateful to Dr. Jeff Kordower for his comments and input, and to members of the Schlossmacher lab for their suggestions.

## FUNDING

This research was funded by: Aligning Science Across Parkinson’s [Grant ID: ASAP-020625] through the Michael J. Fox Foundation for Parkinson’s Research (MJFF); a Canadian Institutes of Health Project Grant; the Uttra and Sam Bhargava Family; the Ottawa Hospital Foundation (to M.G.S.); the Parkinson Research Consortium Ottawa (M.G.S.; J.J.T.); and a uOttawa Brain & Mind Research Institute Team Grant (M.G.S., J.J.T., S.S.). M.L. received support through a CGS-D Award from the Canadian Institutes of Health Research. K.T. was supported by a Francis Matthew Memorial Fellowship from the Parkinson Research Consortium and uOttawa Brain & Mind Research Institute. The authors declare no conflict of interest with the content of this article.

**Supplementary Figure 1. Lrrk2 p.D1994S does not alter survival, weight change, or viral titres in the lungs of mice nasally inoculated with influenza A virus, H1N1.** Survival is graphically displayed as percent survival against days p.i. (**A, D**), and weights are shown as percent of initial body weight against days p.i. (**B, E**). Data are shown separately for female (**A-C**) and male mice (**D-F**). Viral titres were measured in the lungs at 3 days p.i (**C, F**). Levels of H_2_O_2_ are presented for males (**G**) and females (**H**) (measurements shown are relative to brain weight). Each symbol represents one animal. Error bars represent mean ± SEM. WT = wild-type; DS = Lrrk2 p.D1994S mutant mice. Statistical tests used: log rank (Mantel-Cox) test for survival curves; one-way ANOVA for viral titre analyses; two-way ANOVA with uncorrected Fisher’s LSD post-hoc test. *P*-value <0.05 (*), *P*-value<0.01 (**).

**Supplementary Figure 2. Lrrk2 p.G2019S expression alters tau and α-synuclein metabolism as well as oxidative stress in the brains of male mice following H1N1 influenza pneumonitis.** Adult mock-(H1N1 -) and H1N1-infected (H1N1 +) male mice (aged 5-7 weeks) that survived past 14 days post-inoculation were aged for an additional 4 weeks and their brains were collected. Bar graphs indicate levels of total tau protein (A), phosphorylated tau (pSer_199_) (B), the ratio of phosphorylated tau-to total-tau levels (C); total α-synuclein (performed by ELISA; D), phosphorylated α-synuclein (pSer_129_) (E), and the ratio of phosphorylated α-synuclein-to-total α-synuclein (both performed by Western blotting; **F**). Levels of Plk2 (**G**), H_2_O_2_ (**H**), and protein nitrotyrosination content (**I**) of the brain (H_2_O_2_ and nitrotyrosination measurements shown are relative to brain weight). Each symbol represents one animal. Error bars represent mean ± SEM. WT = wild-type; GS = Lrrk2 p.G2019S mutant mice; DS = Lrrk2 p.D1994S mutant mice. Statistical analysis: two-way ANOVA with uncorrected Fisher’s LSD post-hoc test. *P*-value <0.05 (*), *P*-value<0.01 (**).

**Supplementary Figure 3. Lrrk2 p.G2019S expression alters tau and α-synuclein metabolism as well as oxidative stress in the brains of female mice following H1N1 influenza pneumonitis.** Adult mock-(H1N1 -) and H1N1-infected (H1N1 +) female mice (aged 5-7 weeks) that survived past 14 days post-inoculation were aged for an additional 4 weeks and their brains were collected. Bar graphs indicate levels of total tau protein (**A**), phosphorylated tau (pSer_199_) (**B**), the ratio of phosphorylated tau-to total-tau levels (**C**); total α-synuclein (performed by ELISA; **D**), phosphorylated α-synuclein (pSer_129_) (**E**), and the ratio of phosphorylated α-synuclein-to-total α-synuclein (both performed by Western blotting; **F**). Levels of Plk2 (**G**), H_2_O_2_ (**H**), and protein nitrotyrosination content (**I**) of the brain (H_2_O_2_ and nitrotyrosination measurements shown are relative to brain weight). Each symbol represents one animal. Error bars represent mean ± SEM. WT = wild-type; GS = Lrrk2 p.G2019S mutant mice; DS = Lrrk2 p.D1994S mutant mice. Statistical analysis: two-way ANOVA with uncorrected Fisher’s LSD post-hoc test. *P*-value <0.05 (*), *P*-value<0.01 (**), *P*-value <0.001 (***).

## Conflicts of Interest / Financial Disclosures

• Full financial disclosure for the previous 12 months:

Michaela O. Lunn: n/a

Karan Thakur: n/a

Nathalie A. Lengacher: n/a

Quinton Hake-Volling: n/a

Julianna J. Tomlinson: n/a

Earl G. Brown: licensing fees for RNA virus stocks from Roche Pharmaceuticals (2022–2025)

Subash Sad: Canadian Institutes of Health Research; University of Ottawa Brain & Mind Research Institute

Michael G. Schlossmacher: Canadian Institutes of Health Research; University of Ottawa Brain & Mind Research Institute; Brain-Heart-Interconnectome at the University of Ottawa; Aligning Science Across Parkinson’s; Parkinson Research Consortium Ottawa; Jonathan Silverstein Foundation.

