## Supplemental Figures 1-3 for "Elevated phosphorylated α-synuclein and phosphorylated tau in Lrrk2 p.G2019S-mutant mice following influenza A pneumonitis"

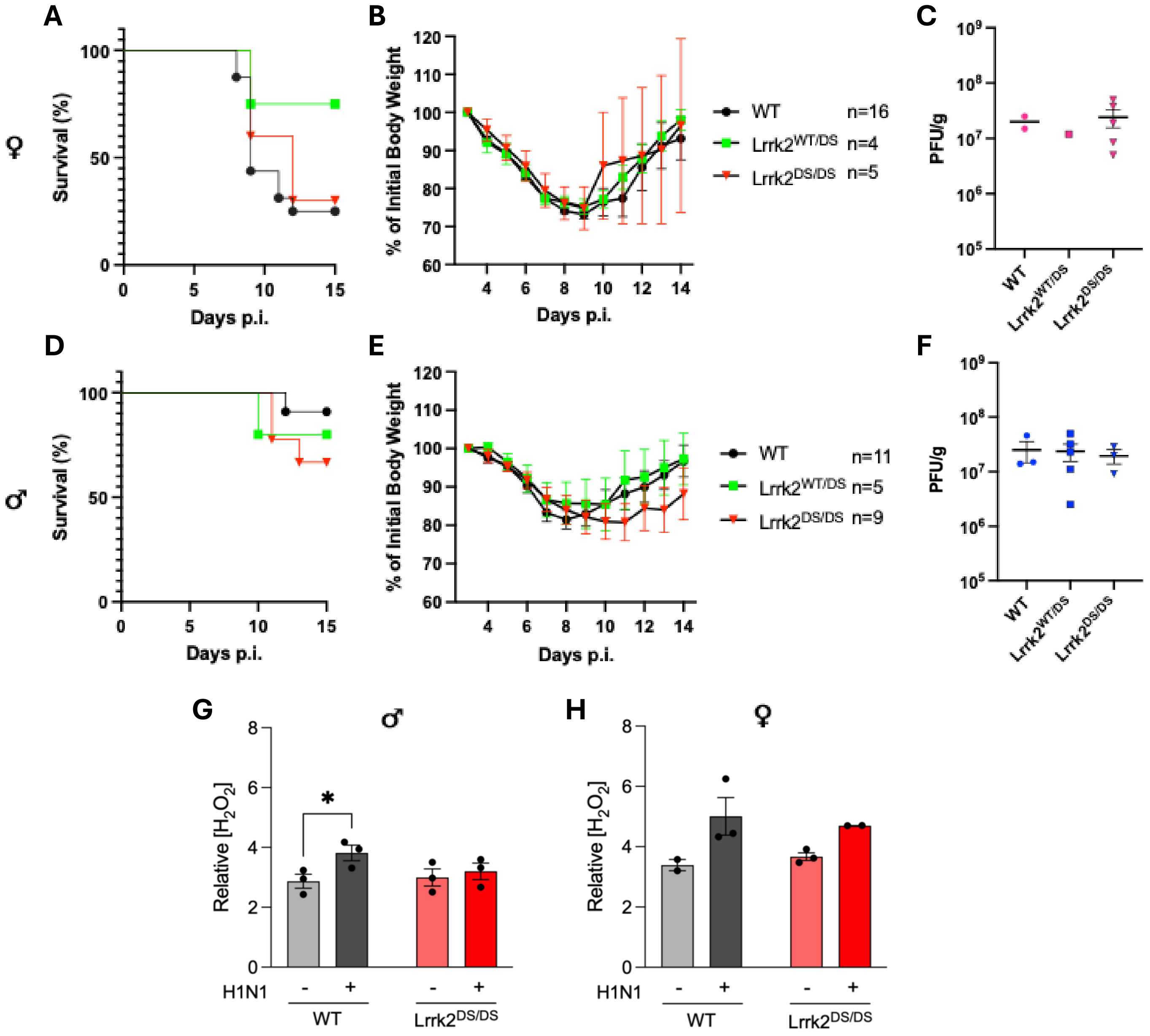


**Supplementary Figure 1. Lrrk2 p.D1994S does not alter survival, weight change, or viral titres in the lungs of mice nasally inoculated with influenza A virus, H1N1**. Survival is graphically displayed as percent survival against days p.i. (**A, D**), and weights are shown as percent of initial body weight against days p.i. (**B, E**). Data are shown separately for female (**A-C**) and male mice (**D-F**). Viral titres were measured in the lungs at 3 days p.i (**C, F**). Levels of $H_2O_2$ are presented for males (**G**) and females (**H**) (measurements shown are relative to brain weight). Each symbol represents one animal. Error bars represent mean ± SEM. WT = wild-type; DS = Lrrk2 p.D1994S mutant mice. Statistical tests used: log rank (Mantel-Cox) test for survival curves; one-way ANOVA for viral titre analyses; two-way ANOVA with uncorrected Fisher's LSD post-hoc test. *P*-value <0.05 (*), *P*-value<0.01 (**).

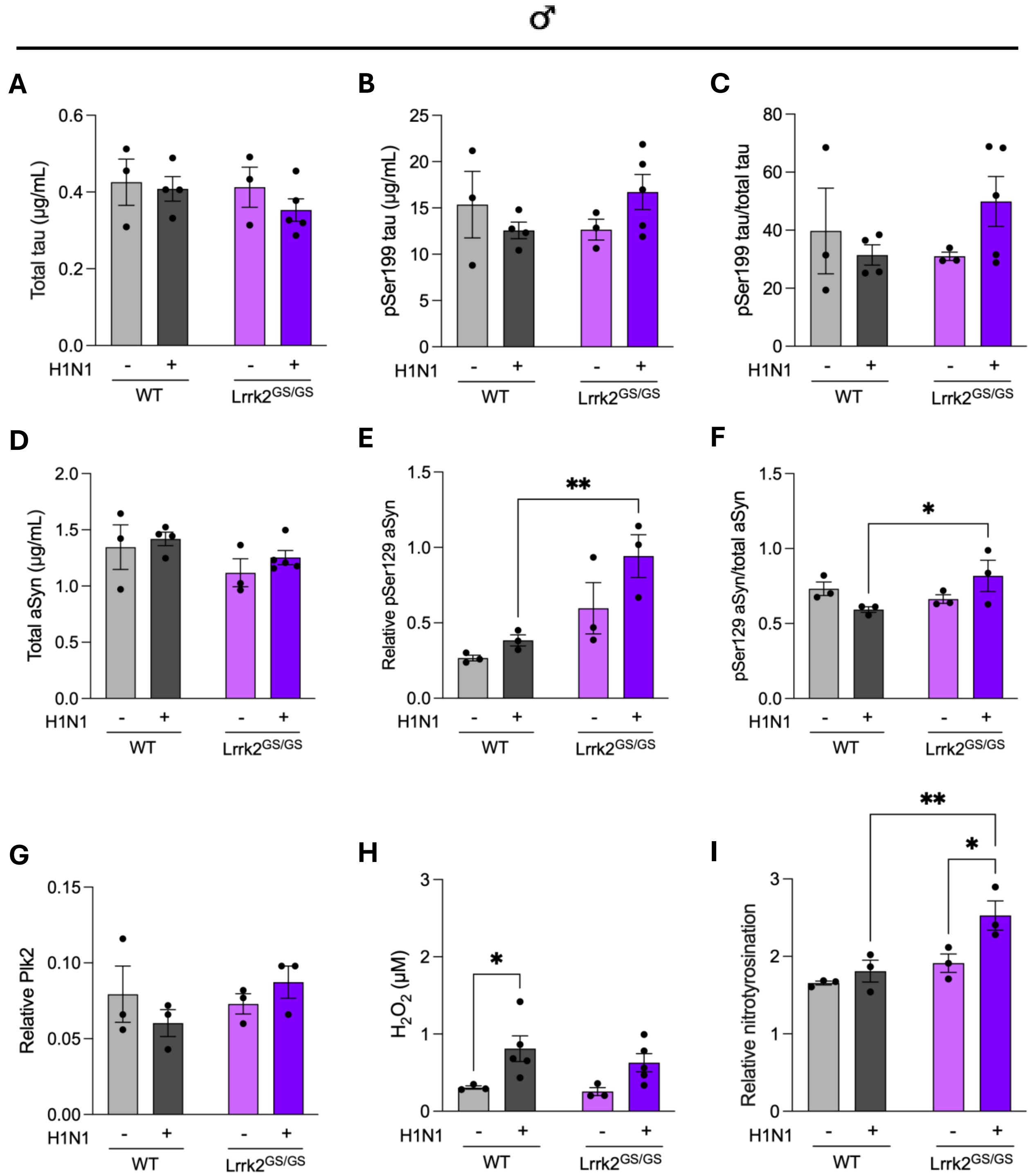


**Supplementary Figure 2. Lrrk2 p.G2019S expression alters tau and α-synuclein metabolism as well as oxidative stress in the brains of male mice following H1N1 influenza pneumonitis.** Adult mock- (H1N1 -) and H1N1-infected (H1N1 +) male mice (aged 5-7 weeks) that survived past 14 days post-inoculation were aged for an additional 4 weeks and their brains were collected. Bar graphs indicate levels of total tau protein (**A**), phosphorylated tau ($pSer_{199}$) (**B**), the ratio of phosphorylated tau-to total-tau levels (**C**); total α-synuclein (performed by ELISA; **D**), phosphorylated α-synuclein ($pSer_{129}$) (**E**), and the ratio of phosphorylated α-synuclein-to-total α-synuclein (both performed by Western blotting; **F**). Levels of Plk2 (**G**), $H_2O_2$ (**H**), and protein nitrotyrosination content (**I**) of the brain ($H_2O_2$ and nitrotyrosination measurements shown are relative to brain weight). Each symbol represents one animal. Error bars represent mean ± SEM. WT = wild-type; GS = Lrrk2 p.G2019S mutant mice; DS = Lrrk2 p.D1994S mutant mice. Statistical analysis: two-way ANOVA with uncorrected Fisher's LSD post-hoc test. *P*-value <0.05 (*), *P*-value<0.01 (**).

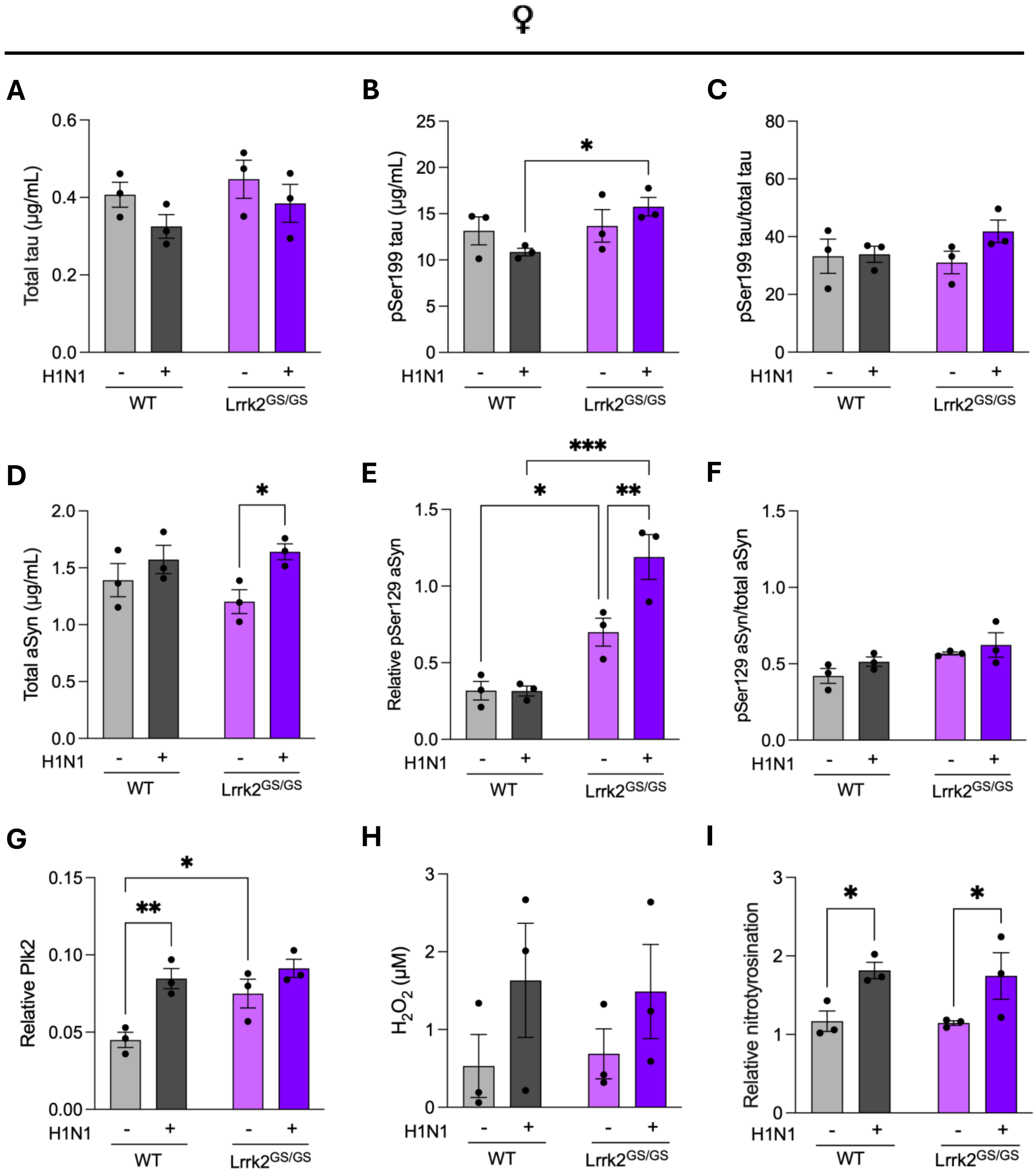


**Supplementary Figure 3. Lrrk2 p.G2019S expression alters tau and α-synuclein metabolism as well as oxidative stress in the brains of female mice following H1N1 influenza pneumonitis.** Adult mock- (H1N1 -) and H1N1-infected (H1N1 +) female mice (aged 5-7 weeks) that survived past 14 days post-inoculation were aged for an additional 4 weeks and their brains were collected. Bar graphs indicate levels of total tau protein (**A**), phosphorylated tau ($pSer_{199}$) (**B**), the ratio of phosphorylated tau-to total-tau levels (**C**); total α-synuclein (performed by ELISA; **D**), phosphorylated α-synuclein ($pSer_{129}$) (**E**), and the ratio of phosphorylated α-synuclein-to-total α-synuclein (both performed by Western blotting; **F**). Levels of Plk2 (**G**), $H_2O_2$ (**H**), and protein nitrotyrosination content (**I**) of the brain ($H_2O_2$ and nitrotyrosination measurements shown are relative to brain weight). Each symbol represents one animal. Error bars represent mean ± SEM. WT = wild-type; GS = Lrrk2 p.G2019S mutant mice; DS = Lrrk2 p.D1994S mutant mice. Statistical analysis: two-way ANOVA with uncorrected Fisher's LSD post-hoc test. *P*-value <0.05 (*), *P*-value<0.01 (**), *P*-value <0.001 (***).
